# Shared and distinct adaptations to early-life exercise training based on inborn fitness

**DOI:** 10.1101/2024.12.04.626895

**Authors:** Daniel G. Sadler, Lillie Treas, Mary Barre, Taylor Ross, James D. Sikes, Ying Zhong, Steven L. Britton, Lauren G. Koch, Umesh Wankhade, Elisabet Børsheim, Craig Porter

## Abstract

**Background:** Low cardiorespiratory fitness due to genetics increases the risk for cardiometabolic disease. Endurance exercise training promotes cardiorespiratory fitness and improves cardiometabolic risk factors, but with great heterogeneity. Here, we tested the hypothesis that the metabolic phenotype imparted by low parental (inborn) cardiorespiratory fitness would be overcome by early-life exercise training, and that exercise adaptations would be influenced in part by inborn fitness.

**Methods:** At 26 days of age, male and female rat low-capacity runners (LCR, *n*=20) and high-capacity runners (HCR, *n*=20) generated by artificial selection were assigned to either sedentary control (CTRL, *n*=10) or voluntary wheel running (VWR, *n*=10) for 6 weeks. Post-intervention, whole-body metabolic phenotyping was performed, and the respiratory function of isolated skeletal muscle and liver mitochondria assayed. Transcriptomics and proteomics were performed on skeletal muscle and liver tissue using RNA-sequencing and mass spectrometry, respectively.

**Results:** Daily VWR volume was 1.8-fold higher in HCR-VWR compared to LCR-VWR. In LCR, VWR reduced adiposity and enhanced glucose tolerance, coincident with elevated total energy expenditure. While intrinsic skeletal muscle mitochondrial respiratory function was unaffected by VWR, estimated skeletal muscle oxidative capacity increased in VWR groups owing to greater mitochondrial content. In the liver, both maximal oxidative capacity and ATP-linked respiration were higher in HCR-VWR than HCR-CTRL. Transcriptomic and proteomic profiling revealed extensive remodeling of skeletal muscle and liver tissue by VWR, elements of which were both shared and distinct based on inborn fitness.

**Summary:** Early-life exercise training partially overcomes the metabolic phenotype imparted by low inborn cardiorespiratory fitness. However, molecular adaptations to VWR are partly influenced by inborn fitness, which may have implications for personalized exercise medicine.

## Introduction

Upward of 60% of youth in the United States are estimated to have low cardiorespiratory fitness (1). Low cardiorespiratory fitness is an established risk factor for all-cause mortality, cardiovascular disease and metabolic syndrome (2–4). Meanwhile, improving cardiorespiratory fitness attenuates cardiovascular disease morbidity and mortality (5) and lessens the risk for type 2 diabetes (6). Genetic factors account for between 50-60% of the population variance in cardiorespiratory fitness (7, 8). The significance of this genetic, or “inborn” component of fitness has been underscored by a rat model (9). Two-way selective breeding of genetically heterogenous rats based upon maximal treadmill running performance produced low- and high-capacity runners (LCR/HCR) with an ∼8-fold difference in exercise capacity (9). Compared to LCR, non-exercise-trained HCR display lessened susceptibility for cardiometabolic diseases, as well as superior mitochondrial respiratory function (10–12).

The remainder of population variance in cardiorespiratory fitness is accounted for by environmental factors, including daily physical activity and structured exercise training. Physical inactivity is linked with low cardiorespiratory fitness and poor health outcomes (13). By contrast, endurance exercise training improves cardiorespiratory fitness and cardiometabolic risk factors (14, 15), potentially mitigating harmful genetic predispositions (16). However, there is profound interindividual variability in the response of both cardiorespiratory fitness and muscle enzyme activity to a standardized dose of exercise training (17, 18). Evidence from human and rodent model studies suggests an important role for genetic factors in explaining this heterogeneity (19–22). Interestingly, one previous study has shown that the transcriptional adaptation of skeletal muscle to 8-weeks of endurance training was influenced by inborn fitness (23). This result implies potential covariation between the genetic factors driving cardiorespiratory fitness and molecular mechanisms governing training adaptation. However, the extent to which inborn cardiorespiratory fitness affects molecular adaptations to exercise training remains underexplored. Addressing this question could support the development of personalized exercise training programs designed to optimize health outcomes.

To this end, the present study sought to test the following hypotheses: 1) early-life exercise training would mitigate the metabolic defects imparted by low inborn cardiorespiratory fitness; 2) molecular adaptations of skeletal muscle and liver to exercise training would be influenced by inborn fitness. Our results reveal that, in the context of low inborn cardiorespiratory fitness, voluntary wheel running (VWR) for 6 weeks reduces adiposity and improves glucose tolerance without affecting muscle or liver mitochondrial respiratory capacity. Transcriptomic and proteomic profiling of skeletal muscle and liver yielded two key findings. First, VWR did not overcome molecular signatures of low inborn fitness, partly because VWR impacted molecular pathways distinct from those driving cardiorespiratory fitness divergence. Second, despite evidence for shared adaptations to exercise training, such as the repression of hepatic pyruvate kinase, there were substantial strain-specific molecular responses to VWR. Therefore, our data suggest that molecular adaptations to endurance exercise training are influenced, at least in part, by inborn cardiorespiratory fitness.

## Methods

### Animals

Rats selectively bred for low and high running capacity were developed at the University of Toledo, Ohio and underwent rapid quarantine at Charles River Laboratories before arrival at Arkansas Children’s Research Institute. Breeding pairs from generation 47 were sent at approximately 16-20 weeks of age. Upon arrival at our facility, rats were group housed (LCR and HCR separately) at 24°C on a standard light cycle (light 7am – 7pm) with ad libitum access to food (ProLab^®^ RMH 1800 [21.1% protein, 13.8% fat and 65.1% carbohydrate, 4.1 kcal/gram], LabDiet, USA) and drinking water. Virgin male and female rats were subsequently paired for breeding, and after confirming pregnancy, male breeders were removed from the cage. Resultant offspring from a total of 16 litters were weaned at 26 days of age. An a priori power analysis was performed based on our previous data (24) and suggested a sample size of *n*=9 per group would provide an actual power of 0.82 to detect a within strain effect of running wheel access on blood glucose levels (with an allocation ratio of 1). We chose to study *n*=10 animals per group (50% male). Approximately 2-3 offspring were randomly selected from each litter based on body mass (*n*=20 per strain, 50% male) for the purposes of the current study, with the remaining pups being humanely euthanized. Both LCR and HCR were randomized to Control (CTRL) or voluntary wheel running (VWR) groups based on body mass, and subsequently individually housed in standard home cages with or without access to running wheels (Tecniplast Activity Systems, Starr Life Sciences Corp, Oakmont, PA, USA) for 6 weeks, as shown in Figure 1A. In rats randomized to VWR, wheel running was recorded in real time using VitalView® Activity software (Starr Life Sciences Corp, Oakmont, PA, USA). At the end of the study, all rats underwent body composition assessments and glucose tolerance testing and were then euthanized in a rising concentration of CO_2_ between zeitgeber time (ZT) 0-2. All animal procedures were approved by the Institutional Animal Care and Use Committee at the University of Arkansas for Medical Sciences.

**Figure 1.**
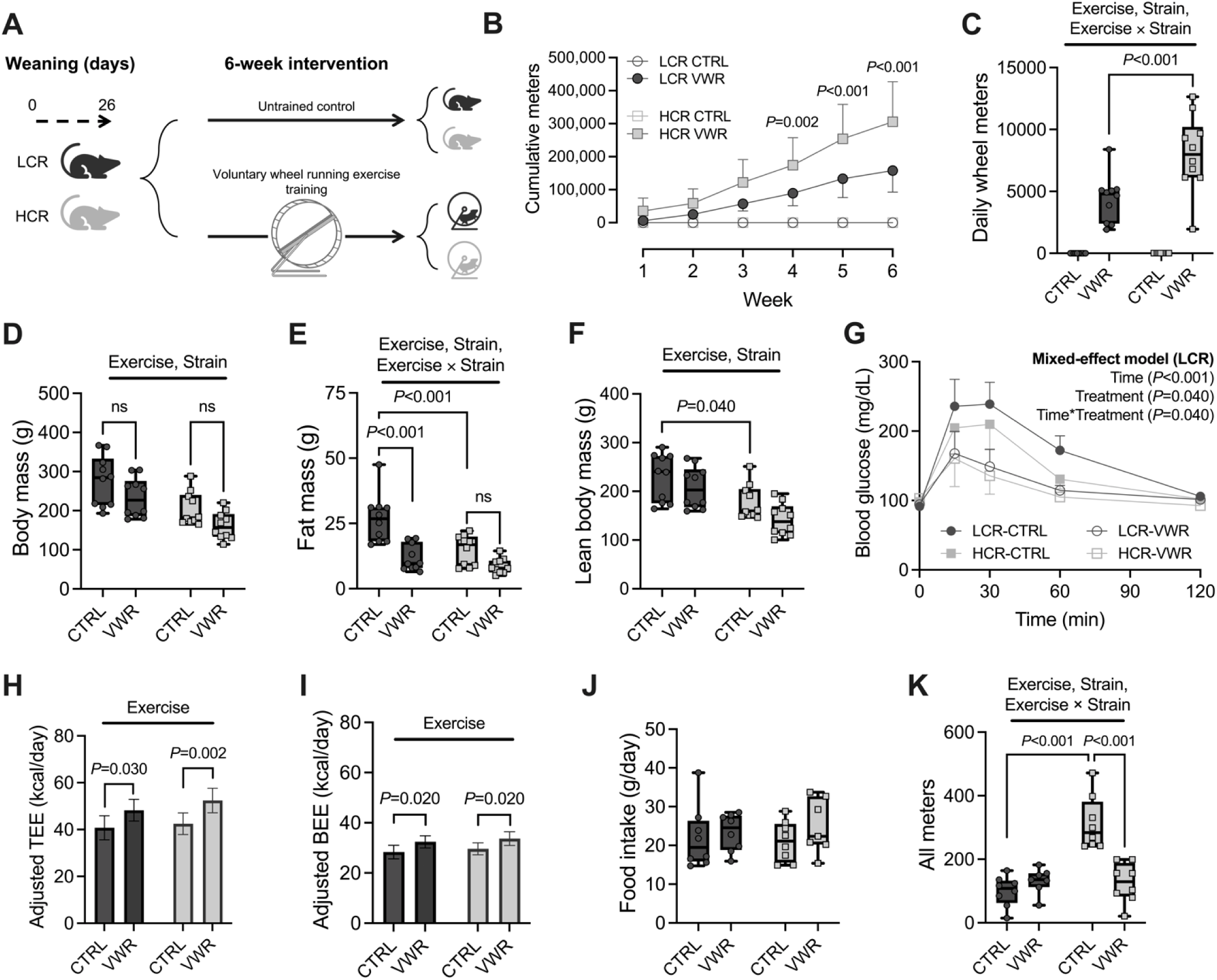
Voluntary wheel running improves metabolic health of juvenile rats with low inborn cardiorespiratory fitness. (A) Experimental design. (B) Cumulative wheel meters ran over the 6-week intervention. (C) Daily wheel meters. (D) Body mass. (E) Fat mass. (F) Lean body mass. *n*=10/group (50% male) for panels A-F. (G) Blood glucose responses to IPGTT. (H) Body mass-adjusted total energy expenditure. (I) Body mass-adjusted basal energy expenditure. (J) Food intake. (K) All meters. *n*=8/group (50% male) for panels G-K. Two-way ANOVA with Tukey’s multiple comparisons test for all panels except G, where mixed-effect model was applied. All data are presented as mean ± SD.

### Body composition and Glucose tolerance

Body composition was measured post-intervention by quantitative magnetic resonance imaging (qMRI) using the EchoMRI-500 (EchoMRI, Houston, Texas, USA). To determine glucose tolerance, rats were injected intraperitoneally with a sterile glucose solution (2 g/kg body mass) after an overnight fast (∼6 pm to ∼ 8 am) and blood samples were taken from the tail at 0, 15-, 30-, 60-, and 120-minutes post injection.

### Metabolic and behavioral phenotyping

Juvenile rats were individually housed in indirect calorimetry cages for 4-5 consecutive days during the final week of the intervention for metabolic and behavioral phenotyping (Sable Systems International, Las Vegas, NV, USA). Rats randomized to VWR were housed with running wheels during metabolic and behavioral phenotyping studies, whereas those randomized to CTRL were not. Rates of oxygen consumption (V O_2_) and carbon dioxide production (V CO_2_) were measured to calculate total energy expenditure (TEE) and its components using the Weir equation (24). Food intake, water intake, locomotor activity and wheel meters were also continuously recorded. Data were analyzed using Sable Systems International ExpeData software (Version 1.9.27). Data from three consecutive 12:12 hour light-dark cycles were averaged to provide daily values.

### Tissue Collection

All animals were euthanized in a rising concentration of CO_2_ prior to tissue collection. For isolation of mitochondria, portions of liver and quadricep tissue were immediately placed in ice-cold mannitol isolation buffer (225 mM mannitol, 75 mM sucrose and 0.2 mM EDTA) or Buffer A (PBS [pH 7.4], supplemented with 10 mM EDTA), respectively. Portions of soleus and liver tissue were also immediately submerged in RNAlater solution (Thermo Fisher Scientific, Waltham, MA) to preserve RNA integrity. Remaining tissue was snap-frozen in liquid nitrogen and stored at -80°C till further analyses.

### Mitochondrial isolation

Mitochondria were isolated from liver and skeletal muscle by two different protocols. Isolation of skeletal muscle mitochondria was based a previously published method (25). Excised quadricep was minced in 1 mL Buffer B (MOPS [50 mM], KCl [100 mM], EGTA [1 mM], MgSO4 [5 mM], pH = 7.1) on ice. The tissue mash was then digested by trypsin (9 mg of 13000-20000 U trypsin/1 g wet mass) for 2.5 minutes with continuous stirring. After digestion, Buffer C (Buffer B containing 2.0 g/L bovine serum albumin) was added to the tissue suspension, before being homogenized with a glass PFTE pestle for 6 strokes. The resulting homogenate was centrifuged for 10 minutes at 800 *g*, 4°C. The resulting supernatant was filtered through cheesecloth, before being centrifuged for 10 minutes at 8000 *g*, 4°C. The resulting supernatant was discarded, and mitochondrial pellets were resuspended in Buffer B before undergoing a final centrifugation for 10 minutes at 8000 *g*, 4°C. Isolation of hepatic mitochondria was based on the protocol by Sumbalova *et al*., (2016). Approximately 500 mg liver was minced in 6 mL mannitol buffer (225 mM mannitol, 75 mM sucrose and 0.2 mM EDTA) on ice and homogenized with a glass PFTE pestle for 5 strokes. The resulting homogenate was brought up to a final volume of 10 mL with mannitol buffer and centrifuged for 10 minutes at 800 *g*, 4 °C. The resulting supernatant was filtered through cheesecloth, before being centrifuged for 10 minutes at 8000 *g*, 4°C. The supernatant was removed, and mitochondrial pellets were resuspended in mannitol buffer and centrifugated for 10 minutes at 8000 *g*, 4°C. Again, the supernatant was removed, and mitochondrial pellets were resuspended in mannitol buffer and centrifugated once more for 10 minutes at 8000 *g*, 4°C. Resulting mitochondrial pellets were resuspended in mannitol buffer and protein contents of final mitochondrial pellets were determined by the BCA assay.

### High-resolution Respirometry

Between 50-100 µg mitochondrial protein was loaded into a 2 mL chamber of an Oxygraph-2K (O2K) high-resolution respirometer (Oroboros Instruments, Innsbruck, Austria) containing 2 mL of MiR05 buffer (MiR05 composition: 0.5 mM EGTA; 3 mM MgCl_2_; 0.5 M K–lactobionate; 20 mM taurine; 10 mM KH_2_PO_4_; 20 mM HEPES; 110 mM sucrose; and 1 mg/ml essential fatty acid free bovine serum albumin) maintained at 37°C. O_2_ concentration was maintained within the range of 30–180 nmol/mL for all analyses, and was recorded at 2–4-s intervals (DatLab, Oroboros Instruments, Innsbruck, Austria). Rates of oxygen consumption (*J*O_2_) were calculated per milligram of mitochondrial protein. Mitochondrial respiration was assayed following the addition of saturating concentrations of complex I-linked substrates (5 mM pyruvate, 2 mM malate and 10 mM glutamate). Thereafter, ADP was added to stimulate maximal coupled respiration and cytochrome c (final 5 µM) was added to establish outer mitochondrial membrane integrity. Succinate was then added to assay complex II-driven respiration before the addition of oligomycin (10 nM final concentration) to quantify ATP-linked respiration. The oxidative capacity of both skeletal muscle and liver tissue was estimated by multiplying respiration linked to ATP synthesis (i.e., oligomycin sensitive respiration per mg mitochondrial protein) by the mitochondrial yield (mg mitochondrial protein per gram of tissue).

### SDS Page and Immunoblotting

Soleus and liver tissue was homogenized in ice-cold 1x precipitation assay buffer containing: 25 mM Tris-HCl pH 7.6, 150 mM NaCl, 1% NP-40, 1% sodium deoxycholate and 0.1% SDS, supplemented with 1x HALT™ protease inhibitor cocktail (ThermoFisher Scientific, USA). Tissue lysates were kept on ice for 1 hour before being centrifuged for 15 minutes at 20,000 × g (4°C). The supernatant was collected and stored at -80°C. Total sample protein was quantified by the Pierce BCA™ assay. Samples were subsequently resuspended in 4x Laemmli buffer (Bio-Rad laboratories, Hertfordshire, UK) containing reducing agent (1x working concentration: 31.5 mM Tris-HCl [pH 6.8], 10% glycerol, 1% SDS, 0.005% Bromophenol Blue and 355 mM 2-mercaptoethanol) and were heated at 95°C for 5 minutes. Between 25-40 µg total protein was loaded based on prior optimization and electrophoresed on 7.5-12.0% Mini-TGX SDS polyacrylamide gels. Proteins were transferred to a PVDF membrane by semi-dry transfer using the Trans-Blot^®^ Turbo™ Transfer System. Ponceau staining was performed after transfer to quantify total lane protein for normalization of protein signals to their respective lanes. After removal of ponceau stain, membranes were blocked in 5% non-fat dry milk for 1 hour at room temperature. After blocking, membranes were incubated overnight at 4°C with rabbit anti total Dusp29 (1:1000; Novus Biologicals, NBP1-84040) or Pklr (1:1000; Novus Biologicals, NBP2-20027) antibodies in bovine serum albumin. Subsequently, membranes were washed 3 times for 5 minutes and incubated for 1 hour in HRP-conjugated anti-rabbit antibody (Cell Signaling Technology, London, UK) at a dilution of 1:20,000 in non-fat dry milk. Proteins were visualised by enhanced chemiluminescence (Thermo Fisher Scientific inc, Waltham, USA) and quantified by densitometry (Amersham Imager 600, GE Healthcare, Life Sciences, NJ, USA).

### RNA extraction and RNA-Seq library preparation

Total RNA was isolated from skeletal muscle and liver tissue using a combination of TRI reagent and RNeasy-mini columns (Qiagen), including on-column DNase digestion. RNA quality and integrity was confirmed by 4200 TapeStation System (Agilent, CA). Equal amounts of total RNA from 2-3 mice were pooled, to generate *n* = 4 biologically distinct replicates per group representing all animals (*n* = 10). Poly-A RNA was isolated from 3 μg of total RNA using Dynabeads® mRNA-Direct kit (Invitrogen). Briefly, poly-A RNA was captured by addition of 30 μL of Oligo-(dt)_25_ Dynabeads in 400 μL of Lysis/Binding buffer. The mixture was incubated on a rotary shaker for 20 min at room temperature. mRNA-bead complexes were washed twice with 400 μL of wash buffer A (10 mM Tris– HCl, pH 7.5, 0.15 M LiCl, 1 mM EDTA, 0.1% LiDS), followed by two washes (400 μL each) with wash buffer B (10 mM Tris–HCl, pH 7.5, 0.15 M LiCl, 1 mM EDTA). mRNA was eluted from the beads in 20 μL of nuclease free water by heating to 65°C for 5 min. The beads were washed twice with wash buffer B (300ul each time), then the second round of mRNA isolation was performed with washed beads and 20ul of mRNA (from first round) in 300 μL of Lysis/Binding buffer. The mixture was incubated on a rotary shaker for 20 min at room temperature. mRNA-bead complexes were washed twice with wash buffer B (300 μL each time). mRNA was eluted from the beads in 11 μL of nuclease free water by heating to 65°C for 5 min. Stranded mRNA-Seq library construction was carried out using 5 μL of purified mRNA (15-75ng) and NEBNext Ultra Directional RNA Library Prep Kit (New England Biolabs). First and second strand cDNA synthesis, purification of double-stranded cDNA using SPRIselect Beads (Beckman Coulter), end repair and dA-tailing, Ligation with NEBNext adaptor (provided in NEBNext oligos kit) was performed with 1:10 diluted adaptor (1.5 µM), in an 83.5 μL reaction volume for 15min at 20°C. Ligated products were purified, and size selected by using SPRIselect Beads. Size-selected cDNA libraries were amplified using indexed primers. PCR was carried out for 11 cycles using 17 μL of template, 2.5 μL of Universal PCR Primer and 2.5 μL of Index(x) Primers (10 μM), and 25 μL of NEBNext Q5 Hot Start HiFi PCR master Mix (New England Biolabs). PCR products were purified using SPRIselect Beads and eluted in 30 μL final volume. A small aliquot (2 μL) was evaluated using High Sensitivity D1000 ScreenTape (Agilent) on a TapeStation 4200 to confirm the absence of primer-dimers and other spurious products. Quantification of the RNA-seq libraries was done via Qubit dsDNA HS Assay kit. Single read 75-bp sequencing of libraries was performed using a NextSeq2000 (Illumina).

### RNA-Seq analysis

FASTQ files were generated following initial quality control with FastQC, adapter trimming, and alignment to the Rattus norvegicus reference genome (Rnor_6.0). Reads with a mapping quality (MAPQ) score greater than 20 were retained, while those below this threshold were filtered out. Trimming and alignment quality control (QC) metrics were evaluated and deemed satisfactory. The resulting BAM files were imported into SeqMonk v1.48.1 for transcript quantification. RNA-Seq quality control plots showed that over 90% of the reads mapped to gene exons. Transcript quantitation was performed using SeqMonk’s built-in RNA-Seq quantitation pipeline, producing log-transformed, normalized counts. These normalized counts were transformed back to raw counts, and differentially expressed genes (DEGs) were identified using the DESeq2 algorithm. To exclude lowly expressed genes that passed the DESeq2 filter, an additional statistical filter based on intensity difference was applied, refining the DEG list to those that were both highly expressed and statistically significant (p < 0.05). The final list of DEGs was generated by combining results from DESeq2 and the intensity difference filter for further analysis.

### Quantitative Proteomics

Total protein from tissue (*n*=10/group/tissue) was reduced, alkylated, and purified by chloroform/methanol extraction prior to digestion with sequencing grade modified porcine trypsin (Promega, Madison, WI, USA). Tryptic peptides were then separated by reverse phase XSelect CSH C18 2.5 um resin (Waters, Milford, MA) on an in-line 150 x 0.075 mm column using an UltiMate 3000 RSLCnano system (Thermo). Peptides were eluted using a 60 min gradient from 98:2 to 65:35 buffer A:B ratio (Buffer A = 0.1% formic acid, 0.5% acetonitrile; Buffer B = 0.1% formic acid, 99.9% acetonitrile). Eluted peptides were ionized by electrospray (2.4kV) followed by mass spectrometric analysis on an Orbitrap Exploris 480 mass spectrometer (Thermo). To assemble a chromatogram library, six gas-phase fractions were acquired on the Orbitrap Exploris with 4 m/z DIA spectra (4 m/z precursor isolation windows at 30,000 resolution, normalized AGC target 100%, maximum inject time 66 ms) using a staggered window pattern from narrow mass ranges using optimized window placements. Precursor spectra were acquired after each DIA duty cycle, spanning the m/z range of the gas-phase fraction (i.e. 496-602 m/z, 60,000 resolution, normalized AGC target 100%, maximum injection time 50 ms). For wide-window acquisitions, the Orbitrap Exploris was configured to acquire a precursor scan (385-1015 m/z, 60,000 resolution, normalized AGC target 100%, maximum injection time 50 ms) followed by 50x 12 m/z DIA spectra (12 m/z precursor isolation windows at 15,000 resolution, normalized AGC target 100%, maximum injection time 33 ms) using a staggered window pattern with optimized window placements. Precursor spectra were acquired after each DIA duty cycle.

### Proteomic Data Analysis

Following acquisition, data were searched using an empirically corrected library and a quantitative analysis was performed to obtain a comprehensive proteomic profile. Proteins were identified and quantified using EncyclopeDIA and visualized with Scaffold DIA using 1% false discovery thresholds at both the protein and peptide level (26). The UniProtKB Rattus norvegicus database was used for the database search. Protein exclusive intensity values were assessed for quality and normalized using ProteiNorm (27). The data was normalized using Cyclic Loess and statistical analysis was performed using Linear Models for Microarray Data (limma) with empirical Bayes (eBayes) smoothing to the standard errors (28). Proteins with an FDR adjusted *P*-value <0.05 and a fold change >1.5 were considered significant. Gene ontology enrichment analysis of proteins significantly different between groups was performed using the online tool Gene Ontology enRIchment anaLysis and visuaLizAtion tool (GOrilla) (29).

### Statistical Analyses

Study data are presented as means ± standard deviation (SD). Two-way ANOVAs were performed using strain and exercise intervention as factors. Pair-wise comparisons were computed using Šídák’s multiple comparisons test. Statistical analyses were carried out using Graphpad Prism version 9 (GraphPad Software, LLC, San Diego, CA, United States). ANCOVA analyses were performed in R Studio (Version 1.4.1717, RStudio, PBC, Boston, MA). Statistical significance was set at *P*<0.05.

## Results

### Early life exercise training improves body composition and glucose tolerance

Over the duration of the 6-week intervention, HCR-VWR accumulated a greater amount of wheel running meters compared with LCR-VWR (Figure 1B). On average, daily wheel running volume of HCR was 1.8-fold higher than LCR (*P*<0.001, Figure 1C). Both strain (Main effect *P*<0.001) and exercise (Main effect *P*=0.009) influenced body mass. LCR-CTRL were heavier than HCR-CTRL (*P*=0.010). VWR tended to decrease body mass relative to CTRL in both strains, although pairwise comparisons did not reach statistical significance (Figure 1D). VWR altered body composition in a strain-dependent manner (Figure 1E-F). Absolute fat mass was higher in LCR-CTRL than HCR-CTRL (*P*<0.001, Figure 1E). VWR reduced absolute fat mass of LCR (*P*<0.001), but not HCR. Absolute lean mass was greater in LCR-CTRL than HCR-CTRL (*P*=0.040, Figure 1F). In LCR, VWR attenuated the accrual of fat mass from pre- to post-intervention, while the accretion of body mass and lean mass were similar between conditions (Supplementary Figure 1A-C). The impact of VWR on glucose tolerance was also strain-dependent. VWR significantly improved glucose tolerance in LCR (Main effect *P*=0.040, Figure 1G), although this effect was absent in HCR. Total energy expenditure was elevated in VWR versus CTRL groups, regardless of strain (both *P*<0.030, Figure 1H). This VWR- mediated increase in total energy expenditure was at least partly attributable to greater activity-related energy expenditure (Exercise main effect *P*=0.001; Supplementary Figure 2A) and increased basal energy expenditure (both *P*=0.020; Figure 1I). The respiratory exchange ratio (Supplementary Figure 2B) and food intake (Figure 1J) were similar between groups. Locomotor activity was influenced by cardiorespiratory fitness (Main effect *P*<0.001) and VWR (Main effect *P*=0.002). HCR- CTRL were more physically active than LCR-CTRL (*P*<0.001, Figure 1K). Meanwhile, VWR attenuated locomotion in HCR rats relative to sedentary HCR (*P*<0.001). Total sedentary time was impacted by strain (Main effect *P*=0.001), whereby HCR-VWR were less sedentary than LCR-VWR (Supplementary Figure 2C).

Relative to body mass, combined mass of the left and right solei was greater in VWR than CTRL groups (Exercise main effect: *P*=0.003, Supplementary Figure 3A). Relative EDL mass was unaltered by VWR but was higher in HCR than LCR (Strain main effect: *P*<0.001, Supplementary Figure 3B). The impact of VWR on relative heart mass depended upon genotype (Interaction: *P*=0.030); heart mass increased significantly with VWR only in HCR (*P*<0.001, Supplementary Figure 3C). A significant main effect of exercise and strain was apparent for relative liver mass, but pair-wise comparisons revealed no significant differences between conditions (Supplementary Figure 3D).

### Early life exercise training does not improve intrinsic mitochondrial respiratory function in rats with low inborn cardiorespiratory fitness

Indices of skeletal muscle mitochondrial respiratory function were not different between any of the groups (Figure 2A). The crude yield of mitochondrial protein from skeletal muscle tissue was comparable between strains, but higher in VWR compared to CTRL groups within each strain (*P*<0.001, Figure 2B). When extrapolating ATP-linked respiration from per gram of mitochondrial protein to per gram of muscle tissue, there was a significant main effect of exercise on estimated ATP-linked respiration (*P*<0.001, Figure 2C). In hepatic mitochondria, complex I-driven leak respiration and maximal respiratory capacity were similar between strains and exercise groups (Figure 2D). However, there was a significant strain and exercise effect on complex I and II-driven hepatic respiratory capacity (*P*=0.002 and *P*=0.030, respectively). VWR improved hepatic respiratory capacity in HCR (*P*=0.040, Figure 2D) but not in LCR. There was also a significant main effect of strain and exercise on ATP-linked respiration (both *P*=0.020). ATP-linked respiration trended higher in HCR-VWR than HCR-CTRL, although this did not reach statistical significance (*P*=0.060). Crude liver mitochondrial yield was similar between conditions (Figure 2E). ATP-linked respiration extrapolated from per gram of mitochondrial protein to per gram of liver tissue revealed a significant main effect of strain on estimated ATP-linked respiration (*P*=0.005, Figure 2F).

**Figure 2.**
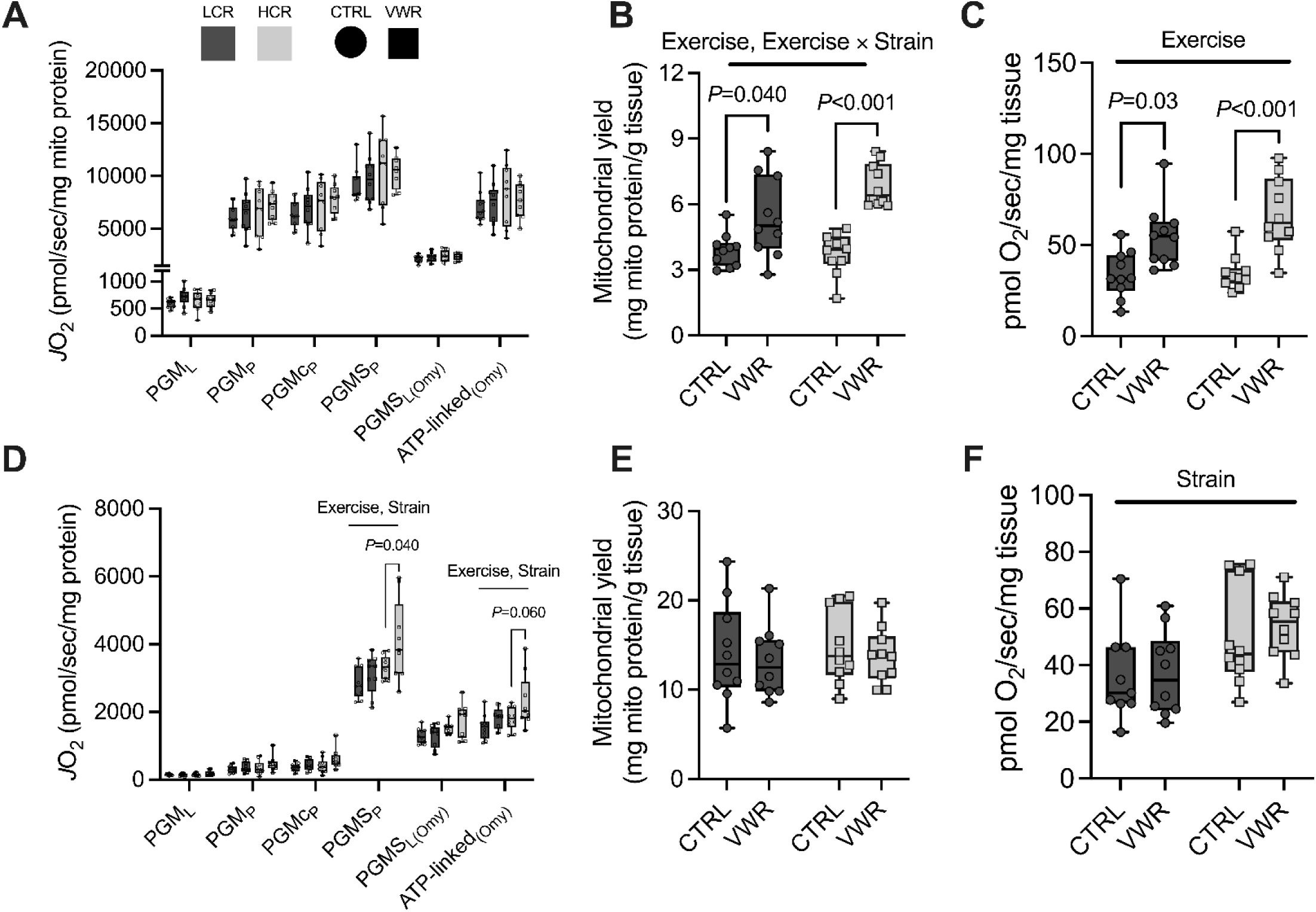
Mitochondrial respiratory function is impacted by voluntary wheel running in a tissue- and strain-dependent manner. Mass-specific rates of oxygen consumption (*J*O_2_) during a substrate, uncoupler and inhibitor titration protocol in isolated (A) skeletal muscle mitochondria and (D) liver mitochondria. Crude mitochondrial protein yields from (B) muscle and (E) liver tissue. Estimated mitochondrial respiration per mg (C) muscle and (F) liver tissue. *n*=10/group. PMGS = Pyruvate (5 mM), malate (2 mM), glutamate (10 mM) and succinate (10 mM). Two-way ANOVA with exercise and strain as main effects, with Sidak multiple comparison test. All data are presented as mean ± SD.

### Transcriptomic and proteomic adaptations to exercise training are influenced in part by inborn fitness

Next, we set out to describe transcriptomic and proteomic responses to VWR across divergent levels of inborn fitness. Before defining the impact of VWR, we compared the molecular signature of LCR and HCR under control conditions. In soleus muscle, 99 DEGs were identified (*q* value < 0.05 and log_2_ FC < 1.5), of which 50 were upregulated and 48 downregulated in HCR-CTRL compared to LCR- CTRL (Figure 3A). DEGs upregulated in HCR muscle were largely enriched in pathways related to lipid metabolism (raw *P* value < 0.05, Figure 3D). Subsequently we defined muscle transcriptional responses to VWR. In total, 142 DEGs were identified in LCR soleus muscle. Of these 142 DEGs, 135 were upregulated and 7 downregulated in LCR-VWR compared to LCR-CTRL (Figure 3B). The upregulated genes were mainly enriched in pathways related to the immune response and cell adhesion (*q* < 0.05, Figure 3E). In HCR muscle, we identified 66 DEGs in response to VWR (56 upregulated and 10 downregulated in HCR-VWR versus HCR-CTRL; Figure 3C). Upregulated genes were associated with various biological processes including the immune response, extracellular matrix organization, cell adhesion and response to hypoxia (*q* < 0.05, Figure 3F). We then asked whether there was evidence for a common transcriptional response to VWR in both strains. We identified 29 genes that were commonly differentially regulated by VWR (28 upregulated and 1 downregulated), accounting for 20% and 44% off all DEGs in LCR and HCR, respectively (Figure 3G). Conversely, 107 genes in LCR and 37 genes in HCR were uniquely differentially regulated by VWR. While the majority of strain-specific DEGs exhibited common directionality in the opposing strain (62-99% of the strain-specific DEGs), they were not significantly impacted by VWR when using only *q* < 0.05 as a significance threshold. Lastly, we asked whether VWR sufficed to rescue the expression of differentially expressed genes in LCR muscle, revealing a single transcript, *Rnf212* (Figure 3H).

**Figure 3.**
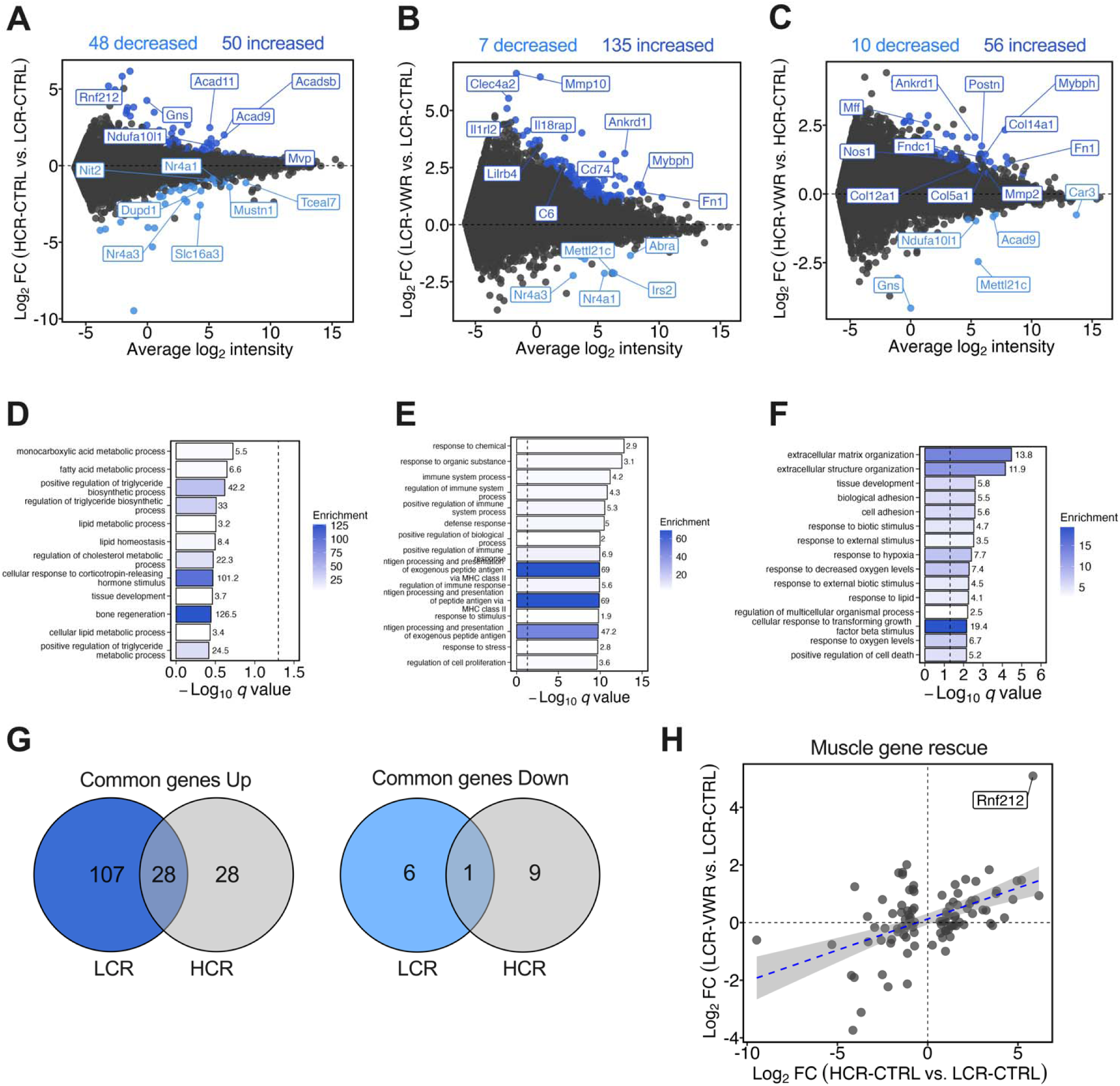
Transcriptomic adaptations of skeletal muscle to exercise training are both shared and distinct based on inborn fitness. MA plots displaying the mean log_2_ intensity versus log_2_ fold change (FC) for transcripts in soleus tissue of (A) HCR-CTRL vs. LCR-CTRL, (B) LCR-VWR vs. LCR-CTRL and (C) HCR-VWR vs. HCR-CTRL. Light and dark blue dots indicate significantly down- or up-regulated genes, respectively. Gene ontology enrichment analysis (biological processes) of transcripts significantly upregulated in (D) HCR-CTRL vs. LCR-CTRL (E) LCR-VWR vs. LCR-CTRL and (F) HCR-VWR vs. HCR-CTRL, where bars are colored and numbered by enrichment value. (F) Venn diagrams displaying the number of muscle transcripts up- or down-regulated in response to VWR in both strains. (H) Scatter plot displaying DEGs between HCR-CTRL and LCR-CTRL, comparing log_2_ fold change in transcript levels of HCR-CTRL vs. LCR-CTRL against LCR-VWR vs. LCR-CTRL. *n*=4 biological replicates per group.

At the proteomic level, we identified 97 differentially abundant proteins in the soleus muscle between strains (fold change > 1.5 and *q* < 0.05); 59 proteins were more abundant and 38 were less abundant in HCR-CTRL compared to LCR-CTRL (Figure 4A). Proteins more abundant in HCR-CTRL muscle were enriched in biological processes such as fatty acid and carboxylic metabolic processes (*q* value < 0.05; Figure 4D). Next, we defined proteomic responses to VWR in both strains. In total, 119 differentially abundant proteins were identified in LCR muscle in response to VWR. Of these 119 proteins, 83 were more abundant and 36 were less abundant in LCR-VWR compared to LCR-CTRL (Figure 4B). Proteins more abundant in LCR-VWR were largely related to cell adhesion processes (*q* value < 0.05, Figure 4E). In total, 37 differentially abundant proteins were identified in HCR with VWR; 14 were more abundant and 23 were less abundant in HCR-VWR compared to HCR-CTRL (Figure 4C). We then asked whether there was a shared muscle proteomic response to VWR in both strains. In sum, we identified 15 commonly differentially abundant muscle proteins (6 increased and 9 decreased) in response to VWR (Figure 4F), encompassing 13% and 41% off all differentially abundant proteins in LCR and HCR, respectively. Proteins identified as differentially abundant uniquely in either LCR or HCR mostly exhibited common directionality in the opposing strain (81- 91%). However, they were still not significantly impacted by VWR when using a less strict significance threshold (*q* < 0.05 only). Lastly, we asked whether VWR was able to normalize the abundance of proteins differentially expressed in LCR muscle. In total, 5 proteins met this criterion, of which two were increased (*Mvp* and *Tapbp* proteins) and 3 were decreased (*Gpd1*, *Tceal7*, and *Pdlim7* proteins) in response to VWR (Figure 4G).

**Figure 4.**
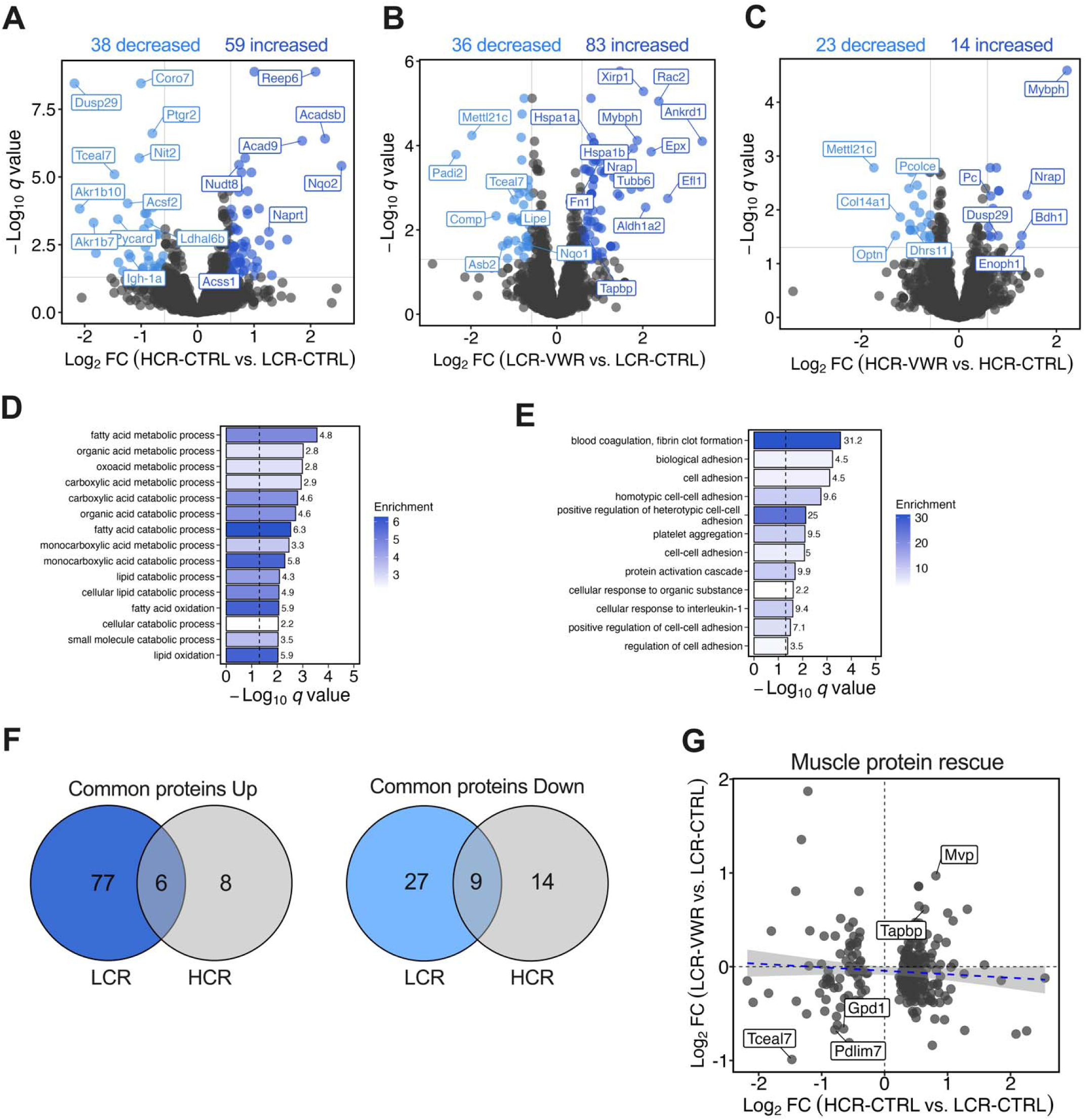
Skeletal muscle proteomic adaptations to endurance exercise training are both shared and distinct based on inborn fitness. Volcano plots comparing the log_2_ fold change in protein abundance against log_10_ *q* values in soleus muscle tissue of (A) HCR-CTRL vs. LCR-CTRL, (B) LCR-VWR vs. LCR-CTRL and (C) HCR-VWR vs. HCR-CTRL. Gene ontology enrichment analysis (biological processes) of proteins more abundant in (D) HCR-CTRL vs. LCR-CTRL and (E) LCR-VWR vs. LCR-CTRL, where bars are colored and numbered by enrichment value. (F) Venn diagrams of muscle proteins either more or less abundant in response to VWR in both strains. (G) Scatter plot displaying differentially abundant proteins between HCR-CTRL and LCR-CTRL, comparing log_2_ fold change in protein abundance of HCR-CTRL vs. LCR-CTRL against LCR-VWR vs. LCR-CTRL. Labelled proteins were “rescued” by VWR in LCR. Proteins are labeled by gene name. *n*=10 per group.

We next deciphered if there was any correspondence between transcriptomic and proteomic signatures of muscle in response to selection for inborn fitness or VWR (Supplementary Figure 4A). Among the genes classified as differentially expressed between HCR-CTRL and LCR-CTRL, we identified 10 whose protein products were differentially abundant with common directionality, including *Mvp, Acadsb, Acad9, Acot2, Dusp29 [Dupd1], Reep6, Serhl2, Acsf2, Nit2* and *Tceal7.* Compared to their respective control groups, we identified 20 DEGs in LCR-VWR (*Aldh1a2, Ankrd1, Arpc1b, Capg, Cd44, Col3a1, Coro1a, Cotl1, F13a1, Fn1, Lcp1, Mettl21c, Mrc1, Mybph, Postn, Rpl3, S100a4, Scrn1, Tnc* and *Tubb6)* and 2 DEGs in HCR-VWR (*Mettl21c* and *Mybph*) whose protein products were also differentially abundant with common directionality.

Subsequently, we investigated the transcriptomic and proteomic responses of liver to selection for inborn fitness and VWR. Under control conditions, 98 DEGs were identified (*q* value < 0.05 and log_2_ FC < 1.5), of which 52 were upregulated and 46 were downregulated in HCR-CTRL compared to LCR-CTRL (Figure 5A). DEGs upregulated in HCR liver were associated with lipid metabolic and monocarboxylic acid metabolic processes (*q* value < 0.05, Figure 5D). In response to VWR, 5 DEGs were identified in LCR liver (4 upregulated and 1 downregulated; Figure 5B). Meanwhile, 45 DEGs were identified in HCR liver in response to VWR (32 upregulated and 13 downregulated in HCR-VWR compared to HCR-CTRL; Figure 5C). The upregulated genes in HCR-VWR were enriched in various processes including amino acid catabolism and negative regulation of intestinal lipid absorption (Figure 5E). There was no common hepatic transcriptional response to VWR (Figure 5F). Though a large portion of strain-specific DEGs exhibited common directionality in the opposing strain (40-71%), they did not meet the threshold for statistical significance (*q* < 0.05 only). Lastly, no dysregulated genes in LCR liver were rescued by VWR (Figure 5G).

**Figure 5.**
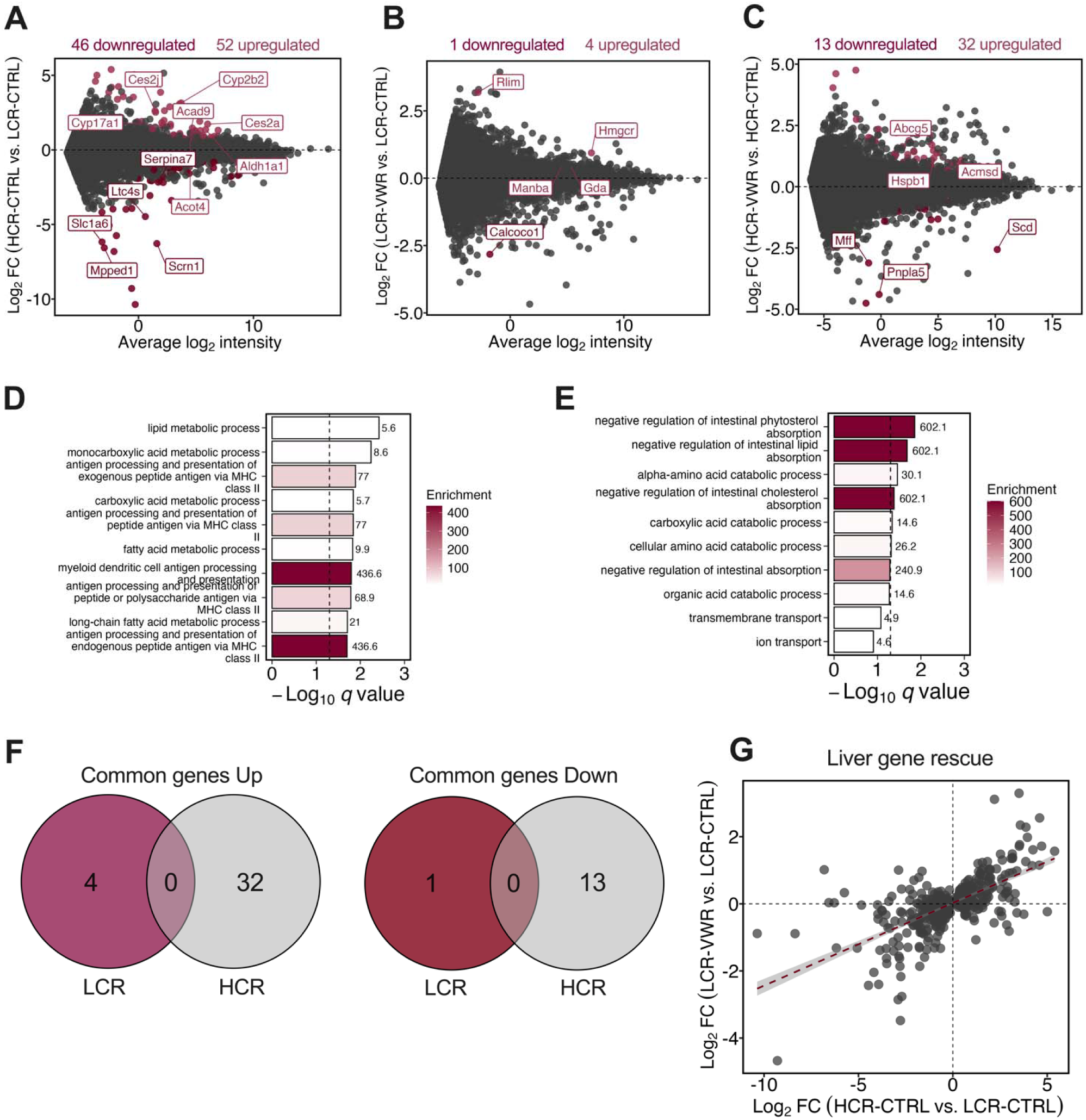
Transcriptomic adaptation of liver in response to selection for inborn fitness and exercise training. MA plots displaying the mean log_2_ intensity versus log_2_ fold change (FC) in transcript abundance in liver tissue of (A) HCR-CTRL vs. LCR-CTRL, (B) LCR-VWR vs. LCR-CTRL and (C) HCR-VWR vs. HCR-CTRL. Light and dark maroon dots indicate significantly up- or down-regulated genes, respectively. Gene ontology enrichment analysis (biological processes) of transcripts significantly upregulated in (D) HCR-CTRL vs. LCR-CTRL and (E) HCR-VWR vs. HCR-CTRL, where bars are colored and numbered by enrichment value. (F) Venn diagrams displaying the number of hepatic transcripts either up- or down-regulated in response to VWR in both strains. (H) Scatter plot displaying DEGs between HCR-CTRL and LCR-CTRL (*q* < 0.05), comparing log_2_ fold change in transcript levels of HCR-CTRL vs. LCR-CTRL against LCR-VWR vs. LCR-CTRL. *n*=4 per group.

In the hepatic proteomic profile, we identified 85 proteins that were differentially abundant between strains, consisting of 50 that were more abundant and 35 that were less abundant in HCR-CTRL compared to LCR-CTRL (Figure 6A). Proteins more abundant in HCR-CTRL liver were enriched in lipid metabolic processes (*q* value < 0.05, Figure 6C). Next, we defined the impact of VWR on the hepatic proteome. A total of 90 differentially abundant liver proteins were identified in LCR in response to VWR (Figure 6B); 80 were more abundant and 10 were less abundant in LCR-VWR versus LCR-CTRL. Hepatic proteins more abundant in LCR-VWR (*q* value < 0.05) were related to cholesterol biosynthesis (Figure 6E). In total, 84 differentially abundant liver proteins were identified in HCR in response to VWR (Figure 6C); 66 were more abundant and 18 were less abundant in HCR- VWR compared to HCR-CTRL. Proteins more abundant in HCR-VWR were enriched in processes related to regulation of endoribonuclease activity, regulation of nucleotide-binding oligomerization domain containing signaling and chaperone-mediated protein folding (raw *P* value < 0.05, Figure 6F). Again, we asked whether there was a shared proteomic response to VWR in both strains. In sum, we identified 33 common differentially abundant liver proteins (31 increased and 2 decreased) in response to VWR with common directionality (Figure 6G), representing 37% and 39% off all differentially abundant proteins in LCR and HCR, respectively. As *Pklr* protein abundance was attenuated in response to VWR in both strains, and both genetic and pharmacological inhibition of this enzyme robustly improves liver health (30), we further corroborated our proteomic data by western blotting (Figure 6H). Analysis of the strain-specific exercise-responsive proteins revealed that 96-100% had comparable directionality in the opposing strain but they were not statistically different (*q* < 0.05 only). We identified 8 dysregulated proteins in the liver of LCR whose abundance was partially rescued by VWR, including *Cyp2e1*, *Cyp3a1*, *Slc26a1*, *Mgst2*, *Tufm*, *Erlin2*, *Rsl1d1l1* and *Pym1* proteins (Figure 6I).

**Figure 6.**
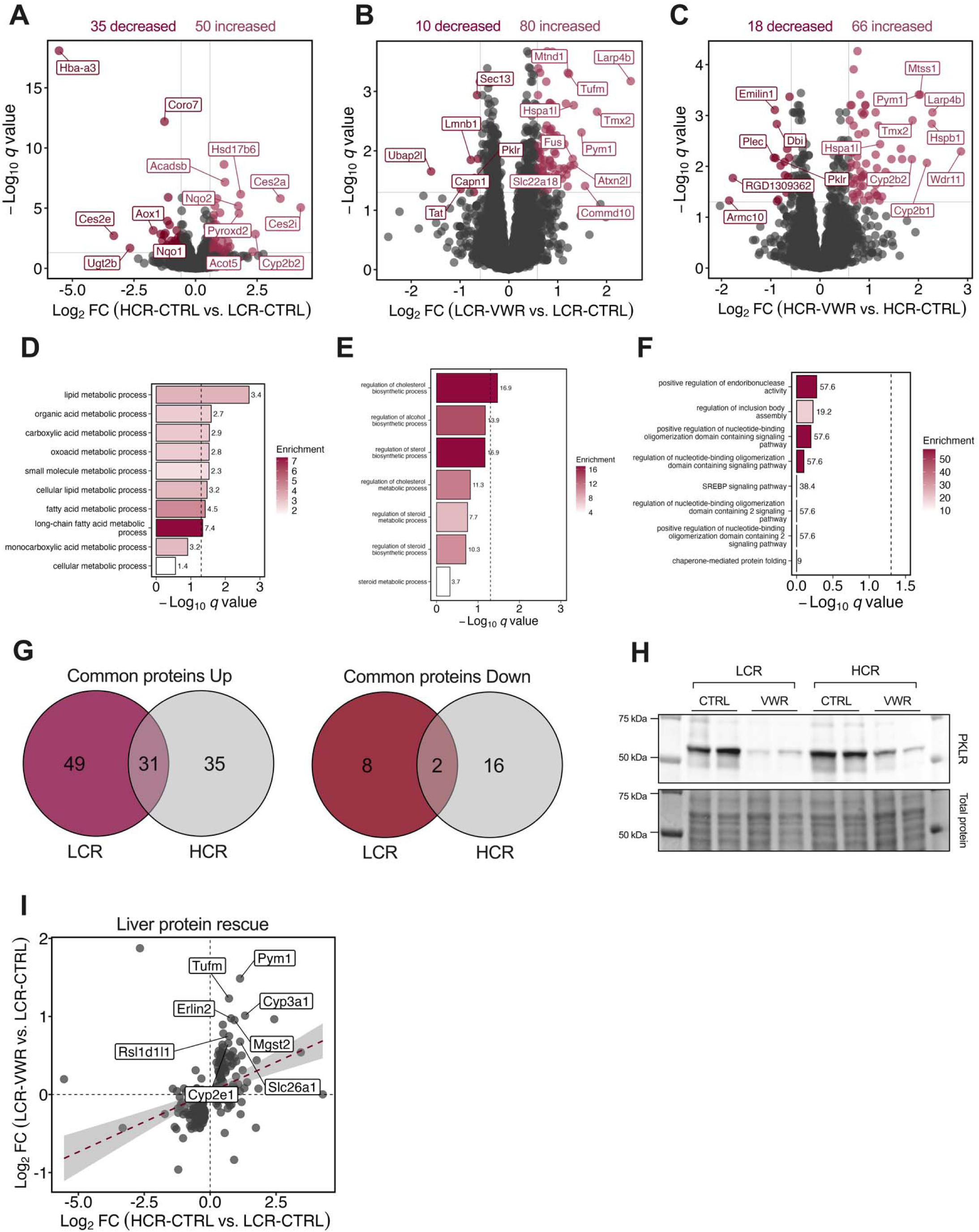
Shared and distinct hepatic proteomic adaptations to exercise training based on inborn cardiorespiratory fitness. Volcano plots comparing the log_2_ fold change in protein abundance against log_10_ q-values in liver tissue of (A) HCR-CTRL vs. LCR-CTRL, (B) LCR-VWR vs. LCR-CTRL and (C) HCR-VWR vs. HCR-CTRL. Gene ontology enrichment analysis (biological processes) of proteins more abundant in (D) HCR-CTRL vs. LCR-CTRL, (E) LCR-VWR vs. LCR-CTRL and (F) HCR-VWR vs. HCR-CTRL, where bars are colored and numbered by enrichment value. (G) Venn diagrams of liver proteins more or less abundant in response to VWR in both strains. (H) Western blot of *Pklr* protein in liver tissue lysates. (I) Liver protein rescue plot displaying all proteins significantly different between HCR-CTRL and LCR-CTRL, with log_2_ fold changes of HCR-CTRL vs. LCR-CTRL on *x*-axis and LCR-VWR vs. LCR-CTRL on *y*-axis. Labelled proteins were “rescued” by VWR in LCR. Proteins are labeled by gene name. n=10 per group.

Finally, we discerned if there was any correspondence between the transcriptomic and proteomic signatures of the liver in response to selection for inborn fitness or VWR. Among the hepatic genes classified as differentially expressed between HCR-CTRL and LCR-CTRL, we identified 9 whose protein products were differentially abundant with common directionality, including *Echdc1, Cyp2b2, Acad9, Anxa5, Acot4, Amdhd1, Aox1, Hba-a3* and *Ces2a.* Compared to their respective control groups, there were no genes in LCR-VWR or HCR-VWR liver whose protein products were differentially abundant with similar directionality (Supplementary Figure 4B).

## Discussion

Exercise-induced adaptations in cardiorespiratory fitness are heterogeneous (17, 31). Much of this variance is owed to genetic factors that are thought to influence the molecular response of tissues to a given exercise stimulus (32, 33). However, there is lack of data concerning the effects of endurance exercise training across divergent levels of inborn cardiorespiratory fitness. To further our understanding of individual responses to exercise training, we sought to investigate whether endurance exercise training mitigates the metabolic phenotype imparted by low inborn cardiorespiratory fitness, and secondly, whether inborn fitness influences molecular adaptations to exercise training. Our results demonstrate that 6 weeks of VWR partially mitigates the metabolic phenotype associated with low inborn cardiorespiratory fitness. Specifically, early-life VWR resulted in lower adiposity and promoted glucose tolerance in LCR, coincident with elevated energy expenditure, while both skeletal muscle and hepatic intrinsic mitochondrial respiratory function were unaffected. Transcriptomic and proteomic profiling of muscle and liver tissue revealed that: 1) the molecular transducers of endurance exercise training may be distinct from those driving divergence in cardiorespiratory fitness; 2) molecular adaptations to exercise training are influenced, at least in part, by genetic predisposition for cardiorespiratory fitness.

### Sedentary LCR exhibit metabolic dysregulation compared to sedentary HCR

It is established that in the absence of exercise training, LCR display greater adiposity, impaired glucose tolerance, and diminished capacity for fatty acid oxidation when compared with HCR (10, 16, 24, 34–36). This phenotypic divergence illustrates the significance of inborn fitness for influencing cardiometabolic disease risk and provides a partial explanation as to why humans with low cardiorespiratory fitness are more likely to have metabolic syndrome and cardiovascular diseases (37, 38). One of the mechanisms thought to be important in driving disparate cardiorespiratory fitness and associated health outcomes between LCR and HCR is the capacity for mitochondrial energy transfer (9). Accordingly, several studies have reported greater mitochondrial content and/or respiratory capacity in permeabilized tissues or isolated mitochondria from HCR compared to LCR rats (11, 12, 35, 39). In this study, we observed no differences in the respiratory capacity of mitochondria isolated from muscle or liver of sedentary LCR and HCR, at least when fueled by complex I- and II-linked substrates. The discrepancy between the present and prior results may be explained in part by the contrasting substrates provided during respirometry, the use of isolated mitochondria versus permeabilized tissue, and/or the muscle origin of isolated mitochondria. Further, animals in this study were still relatively young (∼10 weeks old), whereas most data concerning muscle energetics in HCR versus LCR rats has focused on older adult animals. Indeed, we have previously shown that greater mitochondrial respiratory capacity is more apparent in adult than juvenile HCR rats when compared to age-matched LCR rats (24).

Building on previous research (24, 40, 41), transcriptomic and proteomic profiling of LCR and HCR uncovered molecular disparities that are likely important for driving their phenotypic divergence. A molecular signature reminiscent of impaired lipid catabolism was observed in the skeletal muscle and liver of LCR when compared to HCR, aligning with previous studies (40–43). This manifested as a lower abundance of specific acyl-CoA dehydrogenase and thioesterase enzymes that play essential roles in the oxidation and hydrolysis of fatty acyl-CoA species, respectively. Low lipid oxidative capacity in LCR muscle likely contributes to poor exercise tolerance and may promote metabolic inflexibility (43). While a diminution in hepatic lipid oxidative capacity may underly the susceptibility for diet-induced hepatic steatosis (44). Several other disparities between strains are noteworthy. In soleus muscle, the major vault protein (*Mvp*) was upregulated at the mRNA and protein level in HCR. This protein is a main component of the vault complex – a multi-subunit ribonucleoprotein structure that may act as a scaffold for proteins involved in signal transduction and as a chaperone during nucleo-cytoplasmic transport (45, 46). No prior studies have investigated *Mvp’s* functional role in skeletal muscle. Dual-specificity phosphatase 29 (*Dusp29,* also known as *Dupd1*) was downregulated at the mRNA and protein level in HCR muscle, consistent with prior research (24). This occurred beside downregulation of the transcription factor *Nr4a1*, congruent with a recent study whereby *Dusp29* gene expression highly correlated with an endurance exercise training-mediated decrease in *Nr4a1’s* protein product (47). Overexpression of *Dusp29* in C_2_C_12_ myoblasts was sufficient to reduce ERK1/2 phosphorylation at Tyr 204 (48), but its role in driving phenotypic divergence in cardiorespiratory fitness remains to be elucidated. As for the liver, *Ces2a* gene expression and its protein product were greater in HCR compared to LCR. *Ces2a* is a serine hydrolase highly expressed in the liver. Low hepatic *Ces2* expression and activity have been reported in humans with nonalcoholic steatohepatitis and in human and mouse models of obesity (49, 50). Meanwhile, constitutive whole-body deletion of *Ces2a* in mice leads to the accumulation of diacylglycerol and the exacerbation of glucose intolerance and hepatic steatosis upon feeding of a high-fat diet (51). These findings collectively hint that impaired *Ces2a* expression in LCR may promote fatty liver disease by perturbing diacylglycerol metabolism.

### Physiological and mitochondrial adaptations to exercise training

Established adaptations to endurance exercise training include improved body composition and enhanced glucose homeostasis (52, 53). In this study, VWR promoted a leaner phenotype in LCR, who exhibit greater adiposity than HCR under sedentary conditions. Lower fat mass in LCR rats randomized to VWR was likely supported by greater total daily energy expenditure, which occurred in the absence of altered food intake. Reducing adiposity is critical for promoting glucose tolerance and reducing the risk for the type 2 diabetes in humans with overweight and obesity (54, 55). In line with this, VWR-mediated alterations in adiposity in LCR were associated with modest improvements in glycemic control. These findings suggest that increasing total energy expenditure via physical activity is important for maintaining a healthy body composition and for promoting glucose homeostasis, especially in individuals predisposed to low cardiorespiratory fitness.

Mitochondrial adaptations are also a hallmark of endurance exercise training (56). Here, VWR impacted the respiratory function of isolated mitochondria in a tissue- and strain-dependent fashion. For instance, muscle mitochondrial respiratory function per unit of mitochondrial protein was unaffected by VWR, regardless of inborn cardiorespiratory fitness. Although, estimated muscle oxidative capacity was higher following VWR in both strains when extrapolating respiration from mitochondrial protein to total tissue protein, since VWR resulted in greater mitochondrial protein yields from muscle in both strains, presumably due to exercised-induced mitochondrial biogenesis in skeletal muscle. In contrast, the respiratory capacity and ATP-linked respiration of hepatic mitochondria were enhanced by VWR exclusively in HCR. The apparent strain-dependent effect of VWR on mitochondrial respiration may be owed in part to the disparate exercise training volumes performed by both strains, which could not be controlled for due to the voluntary nature of the exercise paradigm. Another explanation is that genetic variants conserved in selection for low inborn cardiorespiratory fitness resulted in a diminished mitochondrial adaptive response to training in hepatocytes. This premise aligns with recent findings showing that volume-matched high-intensity interval training enhanced the respiratory capacity of interfibrillar muscle mitochondria derived from HCR but not LCR (57). Together, these results demonstrate that VWR does not suffice to augment intrinsic skeletal muscle or liver mitochondrial respiratory function of rats with low inborn cardiorespiratory fitness, that might partly reflect the influence of genetic factors on mitochondrial adaptations to exercise training.

### Exercise training promotes shared and distinct adaptations in peripheral tissues based on inborn fitness

Six weeks of VWR promoted substantial molecular remodeling of skeletal muscle in the present study. This response was seemingly both shared and distinct based on inborn fitness. One common response to VWR we observed was the repression of *Mettl21c* gene expression and its protein product in the muscle of both strains. As a skeletal muscle-specific lysine methyltransferase, *Mett21lc* has been implicated in the regulation of skeletal muscle fiber size by mediating protein degradation (58, 59). Therefore, it is plausible that the VWR-mediated repression of *Mettl21c* aided muscle remodeling by promoting protein degradation. By what mechanism exercise training regulates *Mettl21c* expression and its functional role in muscle adaptation remains an open question. Another interesting result was the apparent ability of VWR to partially rescue the levels of six dysregulated genes or proteins in sedentary LCR muscle, whose functional roles in skeletal muscle exercise adaptation remain largely unknown. One such protein was the product of *Gpd1*, whose abundance was greater in sedentary LCR muscle but reduced by VWR. Consistent with these data, mouse models of *Gpd1* deficiency are associated with improved exercise performance and increased whole-body lipid oxidation (60), but the mechanism by which an exercise-mediated downregulation of *Gpd1* contributes to functional muscle adaptations remains obscure. The observation that VWR sufficed to rescue only a small subset of dysregulated genes or proteins in LCR muscle (and liver for that matter) strongly suggests that distinct mechanisms may underlie skeletal muscle adaptation to aerobic exercise, compared to those driving divergence in cardiorespiratory fitness.

Our understanding of molecular adaptations underlying exercise-induced improvements in liver health is incomplete. The present study revealed that the liver’s proteomic response to VWR was quantitatively more pronounced than its transcriptomic response. This finding is consistent with results from the Molecular Transducers of Physical Activity Consortium animal training study (61) and suggests that post-transcriptional processes may play a crucial role in fine-tuning the global hepatic proteome in response to exercise training. Another key observation was that the liver proteome underwent extensive remodeling in response to VWR, that partly converged in animals with divergent inborn fitness. One common exercise adaptation we observed was the induction of heat-shock proteins, including the protein products of *Hspa1a*, *Hspa1b* and *Hspa1I*. This established response is likely cytoprotective and may help defend against the deleterious effects associated with type 2 diabetes and non-alcoholic fatty liver disease (62–64). We also discovered a previously unreported, common adaptation to exercise training – a reduction in liver *Pklr* protein abundance. *Pklr* protein catalyzes the conversion of phosphoenolpyruvate to pyruvate, the final step in glycolysis. Reduced flux through *Pklr* may limit mitochondrial pyruvate oxidation and liberate phosphoenolpyruvate for gluconeogenesis, supporting increased glucose production during heightened systemic fuel demand. This finding raises two questions: (1) Is repression of hepatic *Pklr* protein necessary for maintaining glucose homeostasis during chronic exercise training? (2) How is *Pklr* expression regulated in response to exercise? The latter may involve the carbohydrate-responsive element-binding protein (*ChREBP*), a transcription factor that binds the *Pklr* promoter (65) and is modulated by glucose and cAMP levels (66). Regardless of the mechanism, exercise-induced repression of *Pklr* protein has important translational implications. Increased *Pklr* expression is associated with more severe hepatic steatosis in humans and mice, while its silencing reduces mitochondrial pyruvate flux, de novo lipogenesis, and steatosis severity in mice (30). In addition to these common exercise adaptations, VWR was sufficient to rescue the levels of several dysregulated hepatic proteins in LCR, emphasizing the potential for regular exercise to mitigate at least some of the deleterious molecular features associated with low inborn fitness.

### Molecular adaptations to exercise training are influenced in part by inborn cardiorespiratory fitness

Our multi-omic analysis revealed that multiple genes and proteins in peripheral tissues were impacted by VWR uniquely in one strain, suggesting that molecular adaptations to exercise training may be influenced by inborn cardiorespiratory fitness. An important caveat is that the average VWR volume was approximately 1.8-fold higher in HCR compared to LCR. Nevertheless, this was unlikely to account for all the heterogeneity in training responses. For instance, the extent of muscle transcriptomic and proteomic remodeling was 2-3 times greater in LCR, even though they engaged in less exercise. Furthermore, our data are consistent with previous research showing that inborn cardiorespiratory fitness affects the transcriptional response of murine skeletal muscle to a standardized dose of treadmill exercise training (23). To our knowledge, this study is the first to report evidence demonstrating that proteomic adaptations to exercise training are influenced partly by genetic predisposition for cardiorespiratory fitness. While prior investigations reported that baseline cardiorespiratory fitness marginally (∼2%) affects trainability of VO_2_ peak in response to 20 weeks of exercise training (67), our results suggest consideration of inborn fitness as an important mediator of the molecular response to chronic exercise training, with potential implications for personalized exercise interventions.

## Summary

Our study demonstrates that early-life VWR partially mitigates the deleterious metabolic phenotype associated with low inborn cardiorespiratory fitness. Six weeks of VWR improved body composition and glycemic control in rats with low inborn fitness but did not fully rescue mitochondrial or molecular deficits. The latter was due in part to the fact that VWR induced pathways dissimilar from those driving divergence in cardiorespiratory fitness. Moreover, our data reveal that the transcriptomic and proteomic responses of skeletal muscle and liver to exercise training are influenced, at least in part, by inborn cardiorespiratory fitness. The presence of both shared and distinct molecular exercise adaptations based on inborn fitness highlights the importance of genetic predisposition in determining exercise training responses. Our findings advocate for the consideration of inborn cardiorespiratory fitness in the optimization of individualized exercise training programs.

## Availability of data and material

All the data presented in this manuscript are available upon request.

## Supporting information

Supplementary Figured 1-4

## Conflict of interest

All authors have no conflict of interest associated with this manuscript.

## Funding

This study was supported by the USDA-ARS (USDA ARS 6026-51000-012-06S). Support was also provided by the NIGMS through P20GM109096 and R35GM142744, and by the Arkansas Children’s Research Institute and the Arkansas Biosciences Institute (ABIPG4622). We acknowledge the IDeA National Resource for Quantitative Proteomics for funding and conducting the proteomic work in described in this study (R24GM137786). The LCR-HCR rat model system was funded by National Institutes of Health Office of Research Infrastructure Programs Grant (P40OD-021331).

## Authors contributions

DGS, CP, EB, SB and LK conceived the study and designed experiments. DGS, LT, MB, JS, YZ, UW and CP collected the data. UW analyzed RNA sequencing data. DGS and CP analyzed and interpreted the rest of the data and wrote the manuscript. All authors reviewed and provided final approval of the version to be published and agree to be accountable for all aspects of the work in ensuring that questions related to the accuracy or integrity of any part of the work are appropriately investigated and resolved. All people designated as authors qualify for authorship, and all those who qualify for authorship are listed. CP is the guarantor for the work and/or conduct of the study, had full access to all the data in the study and takes responsibility for the integrity of data and the accuracy of the data analysis, and controlled the decision to publish.

## Acknowledgements

We acknowledge the technical support of Mr. Trae Pittman and Mr. Bobby Fae. We thank Samantha J. McKee at The University of Toledo for expert care and maintenance of the LCR/HCR rat colony and for aiding with selection of breeder pairs for this study.

## Notes

https://figshare.com/s/326c30d864cb1cdbd80a

### Competing Interest Statement

The authors have declared no competing interest.

### Summary of Updates

References updated in the Discussion section.

