## Supplementary Figured 1-4 for "Shared and distinct adaptations to early-life exercise training based on inborn fitness"

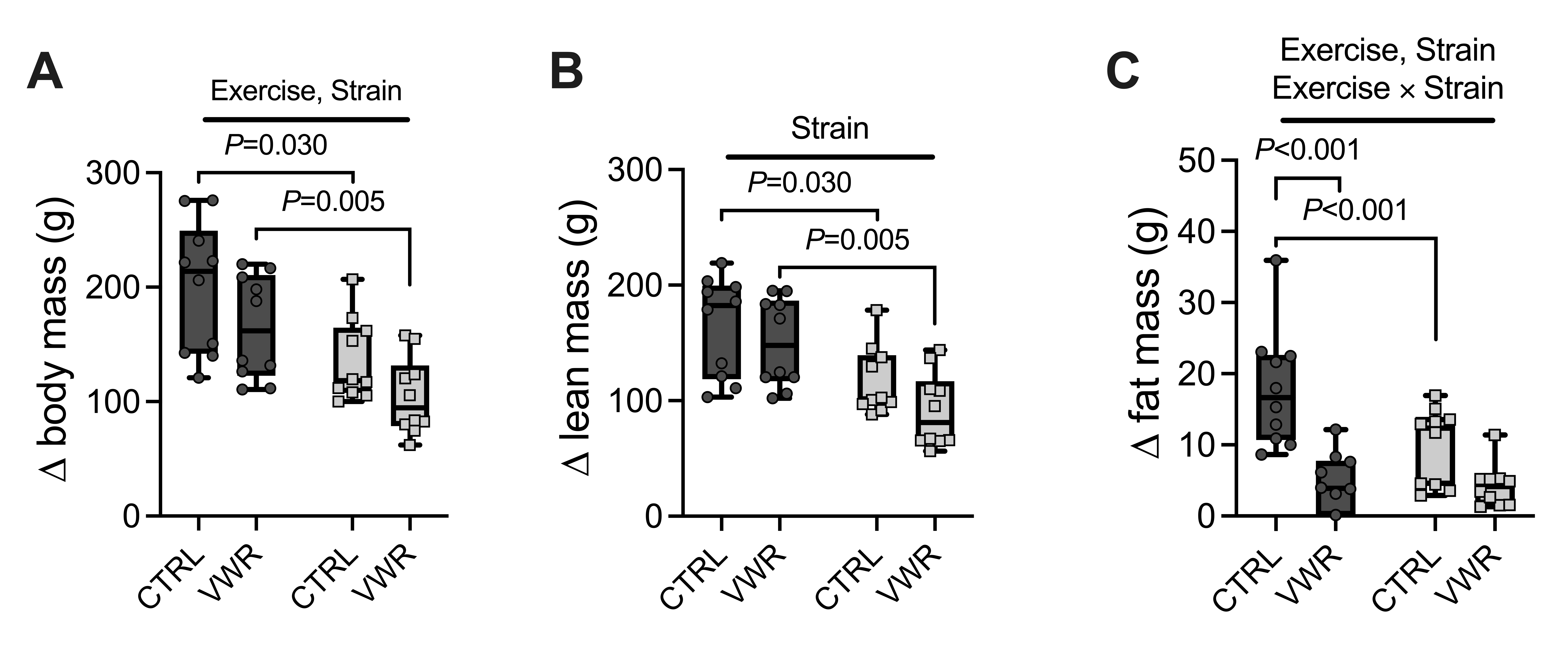

**Supplementary Figure 1.** (A) ∆ body mass. (B) ∆ lean mass. (C) ∆ fat mass. *n*=10/group (50% male). Dark grey and light grey bars/points represent LCR and HCR, respectively. Two-way ANOVA with Tukey’s multiple comparisons test for all panels except G, where mixed-effect model was applied. All data are presented as mean ± SD.

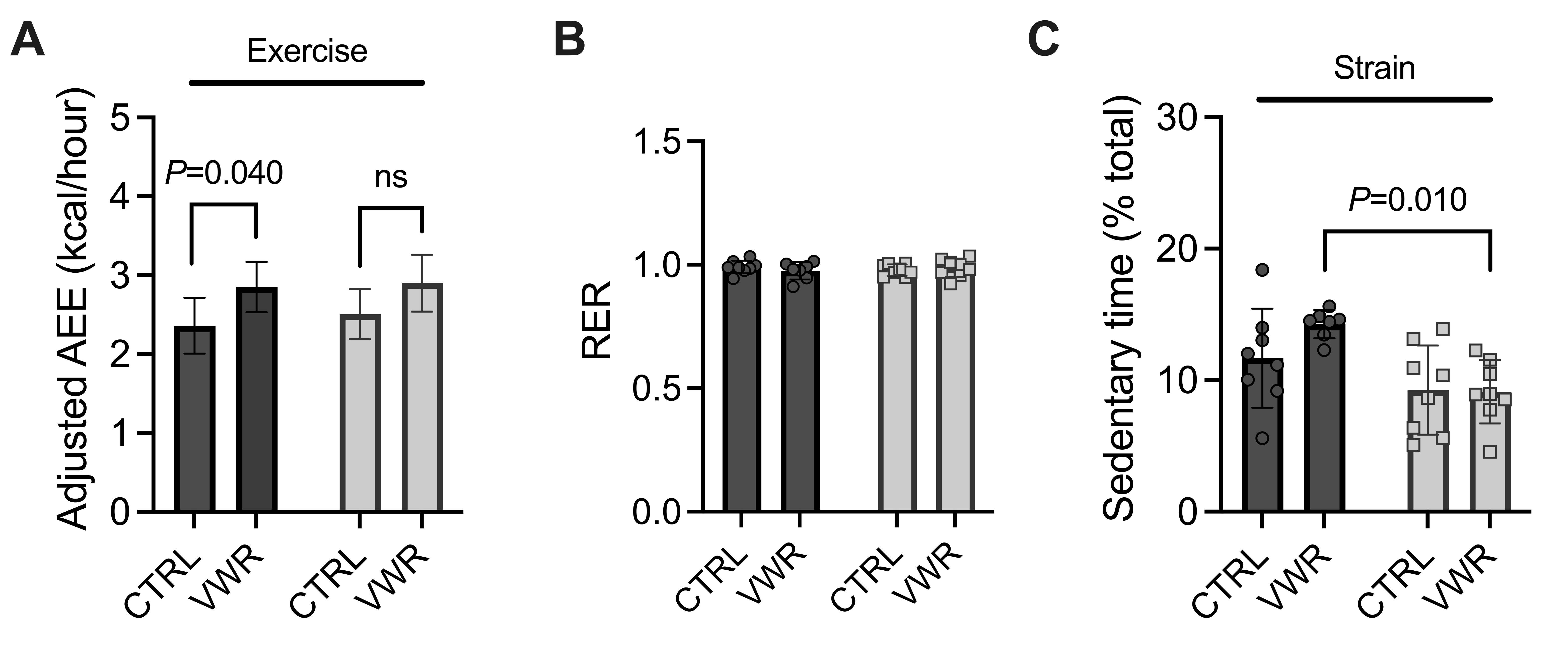

**Supplementary Figure 2.** (A) Adjusted peak activity-related energy expenditure. (B) Respiratory exchange ratio. (C) Daily sedentary time. *n*=8/group (50% male). Dark grey and light grey bars/points represent LCR and HCR, respectively. Two-way ANOVA with Tukey’s multiple comparisons test. All data are presented as mean ± SD.

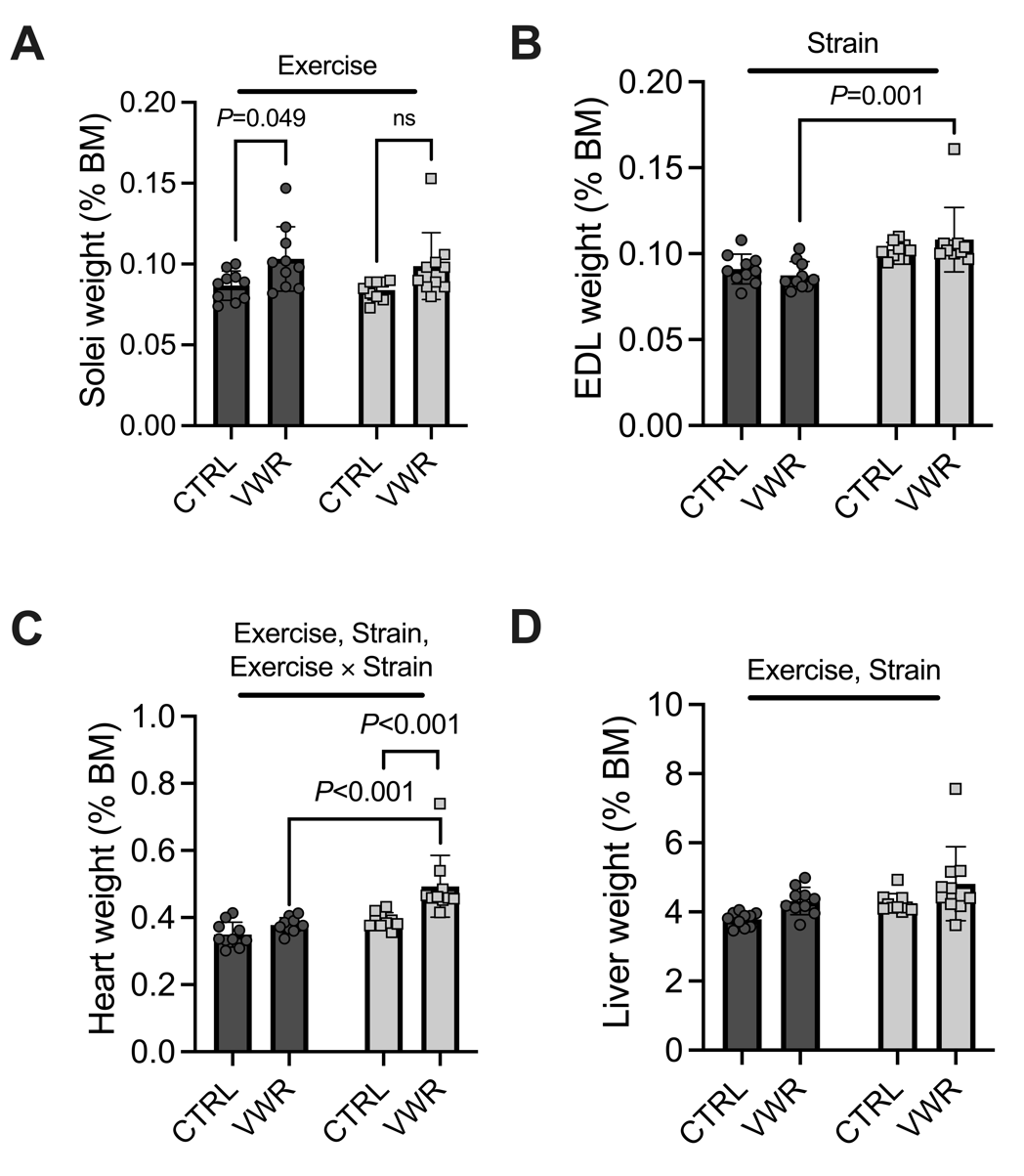

**Supplementary Figure 3.** (A) Solei mass (B) EDL mass. (C) Heart mass (D) Liver mass. Dark grey and light grey bars/points represent LCR and HCR, respectively. Two-way ANOVA with Tukey’s multiple comparisons test. All data are presented as mean ± SD.

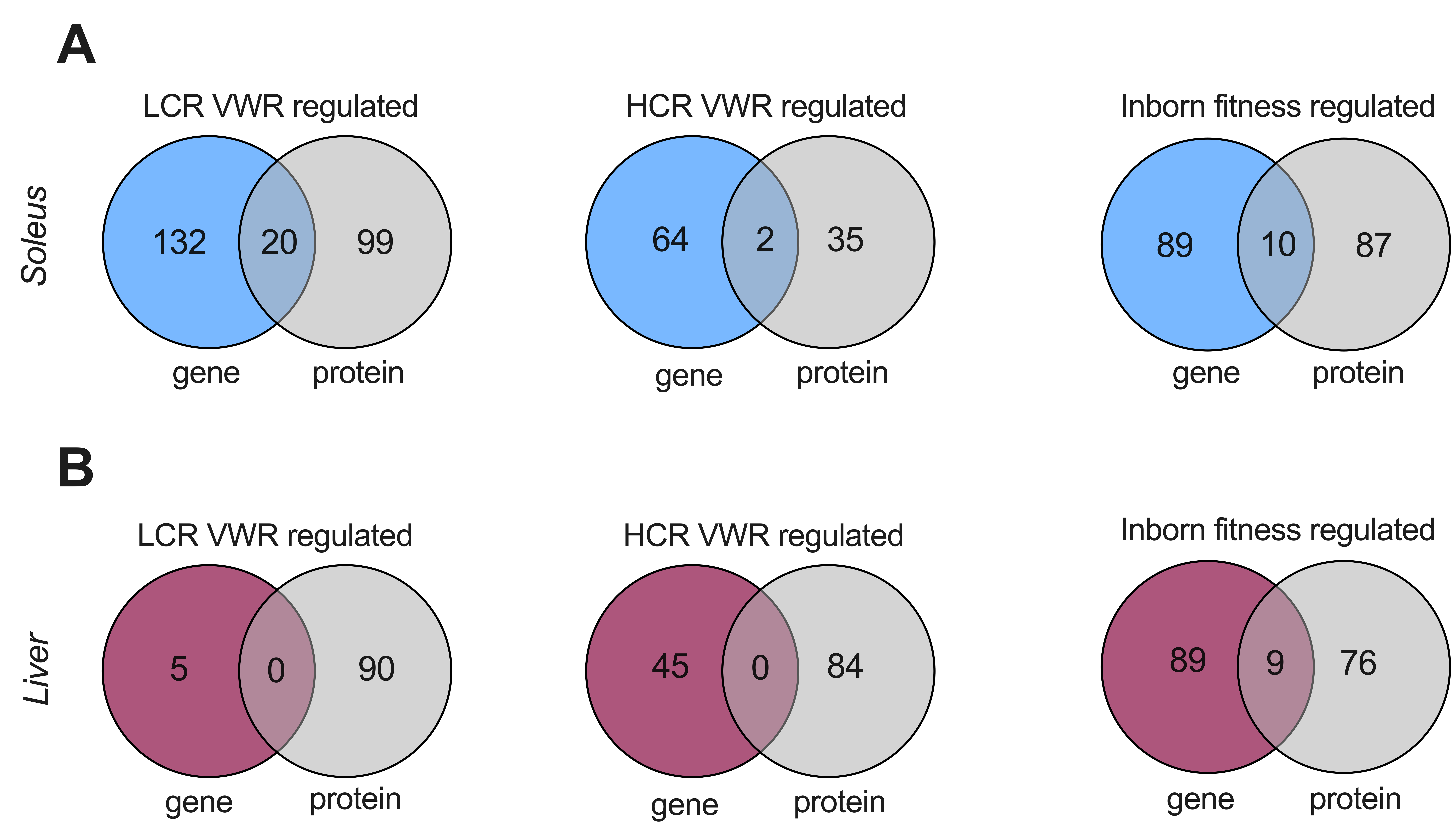

**Supplementary Figure 4.** Venn diagrams of genes and proteins either uniquely or commonly differentially regulated by VWR in LCR and HCR, or by inborn fitness, in (A) soleus muscle and (B) liver.
